# Myotome adaptability confers developmental robustness to somitic myogenesis in response to fibre number alteration

**DOI:** 10.1101/095554

**Authors:** Shukolpa D. Roy, Victoria C. Williams, Tapan G. Pipalia, Kuoyu Li, Christina L. Hammond, Stefanie Knappe, Robert D. Knight, Simon M. Hughes

## Abstract

**Summary Statement:** Homeostatic interactions between muscle stem cells and fibres during myogenesis ensure the correct muscle size is formed independent of fibre number in zebrafish

**Abstract:** Balancing the number of stem cells and their progeny is crucial for tissue development and repair. Here we examine how muscle stem/precursor cell (MPC) numbers are tightly regulated during zebrafish somitic muscle development. MPCs expressing Pax7 are initially located in the dermomyotome (DM) external cell layer, adopt a highly stereotypical distribution and thereafter a proportion of MPCs migrate into the myotome. Regional variations in the proliferation and terminal differentiation of MPCs contribute to growth of the myotome. To probe the robustness of spatiotemporal regulation of MPCs, we compared the behaviour of wild type (wt) MPCs with those in mutant zebrafish that lack the muscle regulatory factor Myod. *Myod^fh261^* mutants form one third fewer multinucleate fast muscle fibres than wt and show a significant expansion of the Pax7^+^ MPC population in the DM. Subsequently, *myod^fh261^* mutant fibres generate more cytoplasm per nucleus, leading to recovery of muscle bulk. In addition, relative to wt siblings, there is an increased number of MPCs in *myod^fh261^* mutants and these migrate prematurely into the myotome, differentiate and contribute to the hypertrophy of existing fibres. Thus, homeostatic reduction of the excess MPCs returns their number to normal levels, but fibre numbers remain low. The GSK3 antagonist BIO prevents MPC migration into the deep myotome, suggesting that canonical Wnt pathway activation maintains the DM in zebrafish, as in amniotes. BIO does not, however, block recovery of the *myod^fh261^* mutant myotome, indicating that homeostasis acts on fibre intrinsic growth to maintain muscle bulk. The findings suggest the existence of a critical window for early fast fibre formation followed by a period in which homeostatic mechanisms regulate myotome growth by controlling fibre size.

## Introduction

How tissue size is regulated is largely unknown, but depends on both the number of cells and their size. When the ‘correct’ size is reached, growth ceases. Although signalling pathways such as IGF, BMP, TOR and Hippo have been implicated in tissue size control (Gokhale and Shingleton, 2015; Irvine and Harvey, 2015), general understanding is lacking. Closely related vertebrate species with distinct ploidy have long been known to alter cell size, yet maintain tissue size through a reduction in cell number (Cavalier-Smith, 2005; Fankhauser, 1945; Gokhale and Shingleton, 2015; Otto, 2007). Thus, tissues appear to measure their absolute size and regulate cell proliferation accordingly, rather than simply generating the correct cell number. Such findings suggest there is feedback regulation between tissue size and stem/precursor cell populations.

Skeletal muscle is a post-mitotic tissue that has the unusual capacity to change size during normal life. All body muscle derives from lineage-restricted stem/precursor cells called myoblasts, that originate from the somitic dermomyotome (Bentzinger et al., 2012). Growth involves three processes: formation of new fibres, fusion of additional myoblasts to existing fibres and increase in cell volume per nucleus. Surprisingly, the contribution of each to tissue growth has not been distinguished in previous studies of embryonic myogenesis. In mammals, fibre formation ceases shortly after birth (Ontell et al., 1988; Ontell and Kozeka, 1984). Fibre number can be a major determinant of muscle size; strains of sheep with different muscle sizes show corresponding differences in fibre number, but not fibre size (Bunger et al., 2009). How myoblasts chose whether to initiate a new fibre or fuse to an existing fibre is unclear. In *Drosophila*, distinct molecular pathways create founder myoblasts, which initiate fibres, and fusion competent myoblasts, which augment fibre growth (Abmayr and Pavlath, 2012). Our recent analyses of zebrafish muscle repair (Knappe et al., 2015; Pipalia et al., 2016) revealed two Pax7-expressing myoblast sub-populations with similarities to founder and fusion-competent cells (Pipalia et al., 2016). Whether such myoblast diversity underlies fibre formation during development and thereby determines muscle size in vertebrates is unknown.

Many genes have been implicated in differentiation and fusion of MPCs marked by Pax7 (Bentzinger et al., 2012). Among them are MyoD and Myogenin, members of the MyoD family of myogenic regulatory transcription factors (MRFs) that drive murine myoblast formation and muscle differentiation (Hasty et al., 1993; Nabeshima et al., 1993; Rawls et al., 1995; Rudnicki et al., 1993; Venuti et al., 1995). MyoD is required for the formation of specific populations of muscle cells early in development, but *Myod* mutants are viable (Kablar et al., 1997; Rudnicki et al., 1992; Tajbakhsh et al., 1997). In contrast, Myogenin appears to be required for differentiation of cells that normally contribute to fusion (Hasty et al., 1993; Nabeshima et al., 1993; Rawls et al., 1995; Venuti et al., 1995). After fibre formation, MRF level within muscle fibres correlates negatively with fibre size and manipulations influence adult fibre size, particularly the response to neurogenic atrophy (Hughes et al., 1999; Moresi et al., 2010; Moretti et al., 2016). Thus, due to their pleiotropic roles, MRFs influence murine muscle size in complex ways

As in amniotes, the zebrafish myotome forms by the terminal differentiation of myoblasts under the control of MRF genes (Hammond et al., 2007; Hinits et al., 2009; Hinits et al., 2011; Maves et al., 2007; Schnapp et al., 2009). In parallel with this process, cells in the anterior somite border generate a Pax3/7-expressing DM external cell layer (Devoto et al., 2006; Groves et al., 2005; Hammond et al., 2007; Hollway et al., 2007; Stellabotte and Devoto, 2007; Stellabotte et al., 2007). Cells of the DM appear to contribute to later muscle growth (Stellabotte et al., 2007). Lineage tracing of zebrafish DM cells suggests that they also contribute to limb and head muscles (Minchin et al., 2013; Neyt et al., 2000). However, quantitative mechanistic understanding of how DM cell dynamics are controlled within the somite and relate to later fibre formation is lacking.

We have previously shown that the zebrafish myotome rapidly increases in volume during the pre- and post-hatching period, growing threefold between 1 and 5 days post-fertilization (dpf) (Hinits et al., 2011). Zebrafish muscle shows size homeostasis in response to altered Myod activity. *Myod* mutants lack specific populations of early myogenic cells so that the myotome is reduced in size by 50% at 1 dpf (Hinits et al., 2009; Hinits et al., 2011). Nevertheless, the myotome of *myod* mutants grows rapidly, approaching normal size by 5 dpf (Hinits et al., 2011). We set out to discover how this happens.

After initial fibre formation in normal growth, dermomyotome-derived Pax7-expressing myogenic cells ingress into the deep myotome around 3 dpf, where a portion express Myogenin and differentiate into fibres, leading to a small increase in fibre number. *Myod* mutants have fewer fibres and fibre number fails to increase. Nevertheless, the remaining fibres grow larger than those in wt. Ingression of Pax7^+^ cells into the myotome is accelerated in *myod* mutants and more cells appear to differentiate. Inhibition of GSK3 activity prevents Pax7^+^ cell ingression, but does not diminish muscle size recovery in *myod* mutants or block growth. The myotome thus responds to reduction in fibre number by hypertrophy of remaining fibres. Our data show that feedback between muscle fibres and their precursor cells regulates myotome growth and that although homeostasis in young animals recovers muscle mass it leaves a persistent alteration in fibre number.

## Results

### Growth of zebrafish muscle

Muscle fibre cross sectional area was determined in embryonic, larval and adult zebrafish (Fig. 1). Mean fibre size increased dramatically in the embryonic period, less rapidly during larval life, slowly beyond 5 months and appeared to plateau after 1 year of age (Fig. 1A). In adults, fibre types were distinguished by myosin heavy chain (MyHC) content (Fig. 1B). As reported previously (Patterson et al., 2008), slow fibres were smaller than the adjacent intermediate fibres, with the more numerous fast fibres in the deep myotome being the largest (Fig. 1A). Paralleling the rapid increase in fibre size from 1-5 dpf, somites increase in mediolateral width (Fig. 1C; p=0.011). Growth also involves increase in fibre number (Fig. 1D). The smallest fibres in developing somites are located near the DM, particularly at the epaxial and hypaxial somitic extremes, suggesting that new fibres arise from DM cells (Stellabotte et al., 2007). Although fibre number increases between 1-3 dpf (Fig. 1D and see below), mean fibre size doubles, despite the lowering effect on mean fibre size of small newly-added fibres (Fig. 1A,D). As around five new slow fibres are formed between 1-3 dpf (Barresi et al 2001), the remaining new fibres must be fast. Counts of nuclei within the myotome also show a 20% increase (Fig. 1D). The increase in myotome nuclei is sufficient to yield five mononucleate slow fibres and twenty extra fast fibres, but does not double like fibre size (Fig. 1D). As shown below, these trends continued until at least 6 dpf. Thus, both increase in fibre volume per nucleus and addition of nuclei to fibres by precursor myoblasts contribute to myotome growth.

**Fig. 1.**
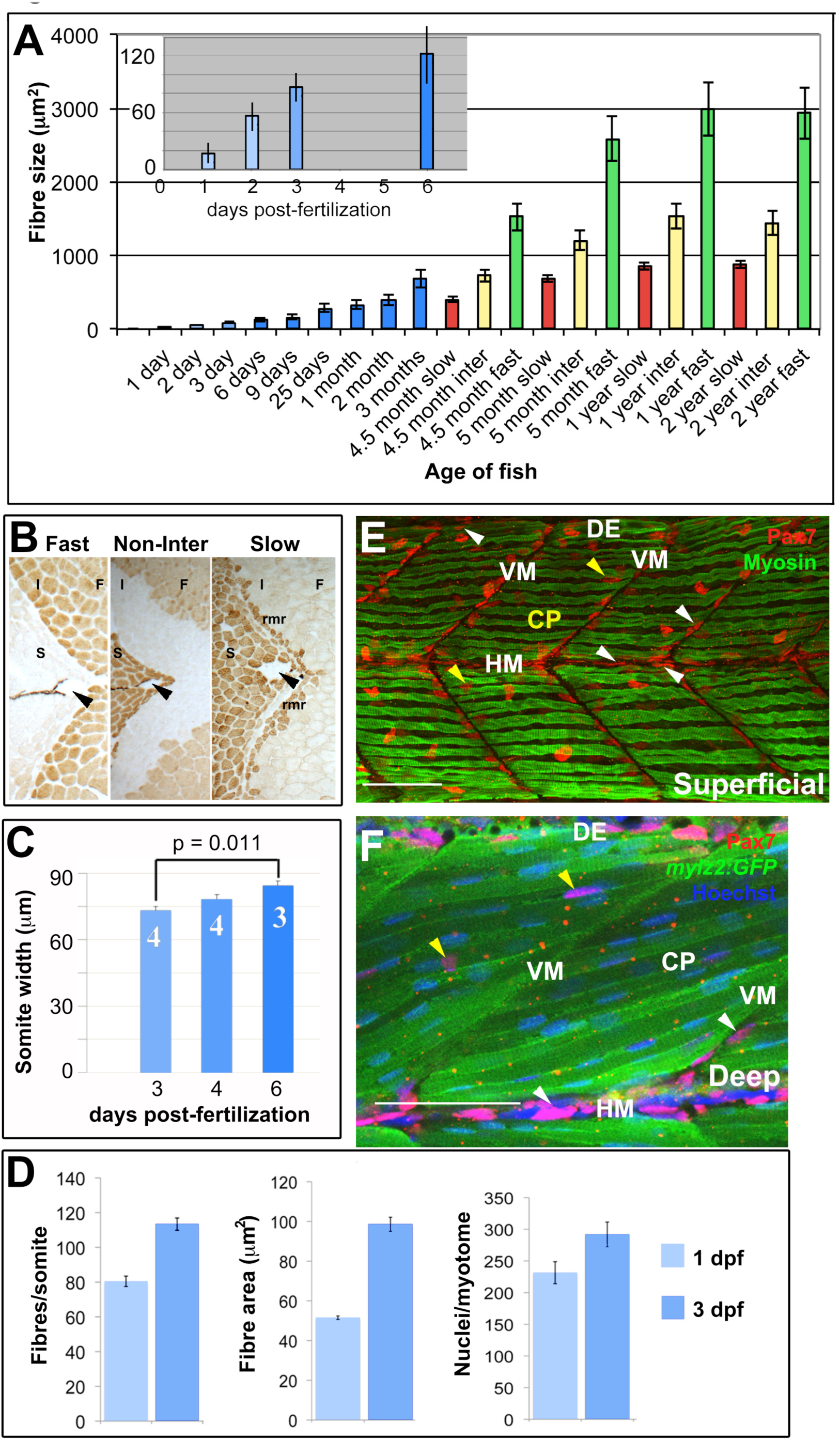
**Muscle growth in relation to Pax7+ cell distribution A.** Fibre cross sectional area from unfixed cryosections as a function of age and fibre type showing all (blue), slow (red), intermediate (yellow) and fast (green) fibres. **B.** Immunodetection distinguishes fast (F), intermediate (I) and slow (S) fibres including the red muscle rim (rmr). Arrowheads indicate the lateral line. **C.** Mediolateral width of somite measured at horizontal myoseptum from wholemount confocal stacks, as in E, F. **D.** Fibres and nuclei were counted and cross sectional area measured on *YZ* confocal sections of somite 16-20 from 8 and 18 lightly-fixed Hoechst-stained *Tg(Ola.Actb:Hsa.HRAS-EGFP)* embryos at 1 and 3 dpf, respectively. As small fibres are hard to count with confidence in fixed preparations, fibre numbers represent minimal estimates. **E, F.** Single confocal slices from wholemount 4 dpf larvae taken in lateral view, orientated with dorsal to top and anterior to left. Wt (E) or *Tg(-2.2mylz2:GFP)^gz8^* (F) larvae stained with anti-Pax7, Hoechst 33342 (detecting nuclei) and either A4.1025 (E, detecting sarcomeric MyHC) or anti-GFP (F). The superficial monolayer of slow muscle fibres aligned parallel to the horizontal myoseptum (HM) in somites 15-18 (E). Pax7^+^ nuclei surround the myotome (white arrowheads) at dorsal edge (DE), HM and vertical myoseptum (VM) and also occur in central portion (CP; yellow arrowheads) in both the epaxial and hypaxial domains. Pax7^+^ cells nestle amongst deeper fast fibres orientated oblique to HM in the epaxial somite (F). Bars 50 μm.

### Increase in Pax7^+^ cells parallels growth of early larval muscle

From where do the extra nuclei and fibres derive? Pax3/7 proteins mark most dermomyotomal MPCs in both amniotes and zebrafish (Devoto et al., 2006; Relaix and Zammit, 2012). Pax7^+^ cells in somites of developing 1 dpf zebrafish are mainly located superficial to differentiated muscle in the DM (Devoto et al., 2006; Feng et al., 2006; Hollway et al., 2007); Fig. 1E,F). New fibres originate at the dorsal edge (DE) of the myotome (Barresi et al., 2001), where many Pax7^+^ cells are located (Fig. 1E,F). As somites mature, Pax7^+^ nuclei accumulate at the vertical and horizontal myosepta (Fig. 1E,F). Pax7^+^ cells at the vertical myoseptum (VM) are initially superficial, near the epidermis, whereas those at the horizontal myoseptum (HM) can be deep within the somite (Fig. 1F). By 5 dpf, small numbers of Pax7^+^ cells are also observed deep within the central portion (CP) of both epaxial and hypaxial myotomes (Fig. 1F). As particular regions of the amniote dermomyotome give rise to distinct MPCs (Buckingham and Rigby, 2014), we analysed the changing numbers of Pax7^+^ cells in defined somitic zones (Fig. S1). Immunolabelling of Pax7 in larvae prior to 4 dpf revealed that most Pax7^+^ cells were located at myotome borders (DE,HM,VM), with the remainder in the superficial CP, the central DM (Figs 1E and 2A). Subsequently, Pax7^+^ cells appeared deep within the somite (Fig. 2A). As no temporal differences in Pax7^+^ cell behaviour in the epaxial and hypaxial somite were noted at any stage examined, and as numbers of Pax7^+^ cells per epaxial somite did not vary detectably along the rostrocaudal axis from somites 14-22 (Fig. S2), we chose to explore changes in Pax7^+^ cell number in the epaxial domain of somites 15-20. Between 3 and 6 dpf, the total number of Pax7^+^ cells per epaxial somite increased by about 50%, from ~40 to ~60 cells (Fig. 2B; p = 0.003). Strikingly, Pax7^+^ cell numbers did not change significantly in the superficial DM; the increase in Pax7^+^ nuclei was accounted for by a rise deep within the somite (Fig. 2A,B; p < 0.001).

**Fig. 2.**
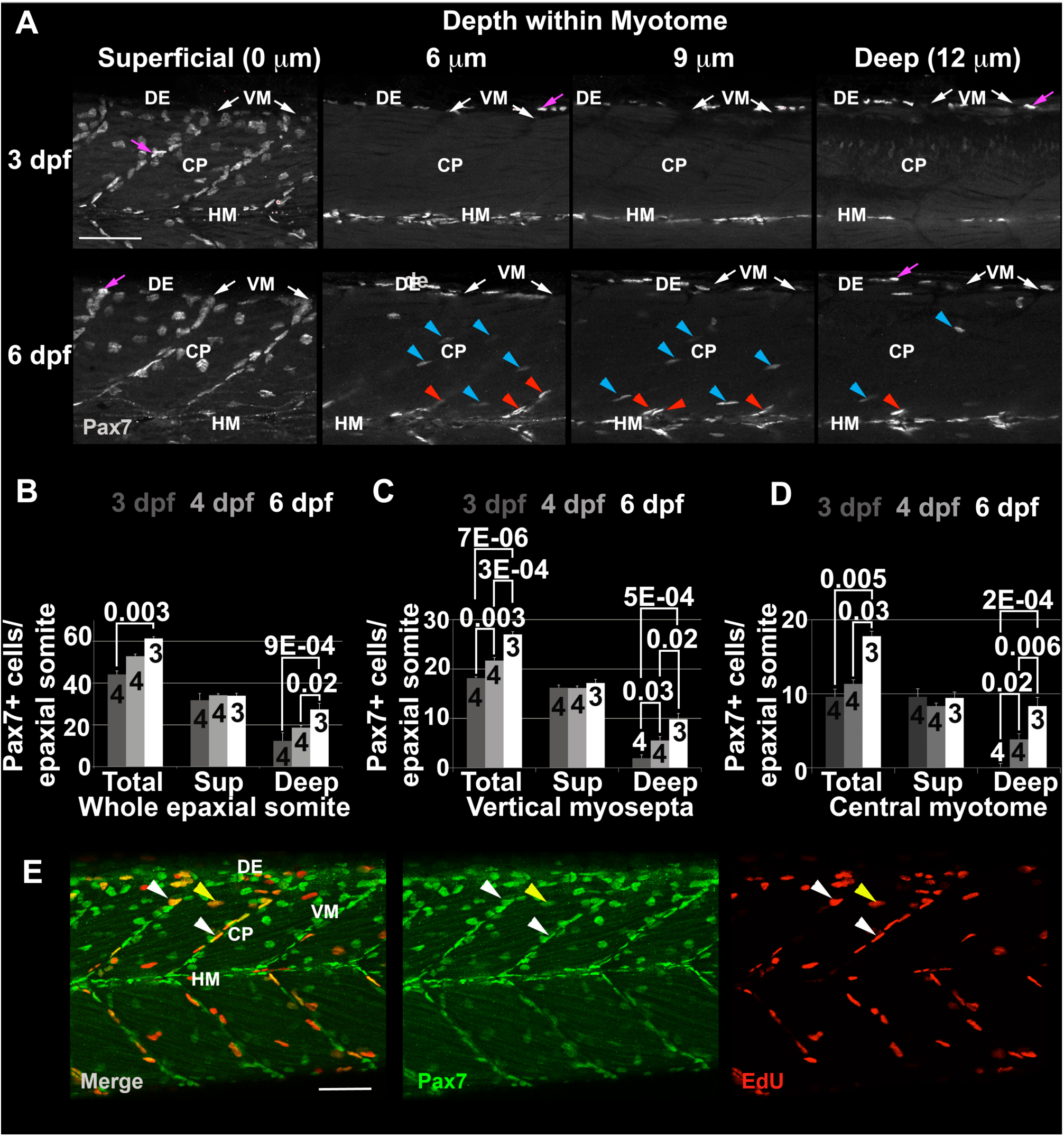
**Pax7^+^ nuclei increase in deep myotome**. Pax7 (A-E) and EdU (E) labelling in wholemount wt larvae. Single confocal slices of zebrafish larvae in lateral view. **A:** Flattened dehydrated embryos imaged at the indicated depths (full larval thickness approximately 35 μm). At 3 dpf, MPCs accumulate superficially near VM (white arrows) but are absent deeper within myotome. Xanthophores (purple arrows) are rare and bright. By 6 dpf, Pax7^+^ nuclei appear deep at the VM (red arrowheads) and CP (blue arrowheads). **B-D:** Numbers of Pax7^+^ nuclei in epaxial somites 16-18 of whole mount larvae increase with age. Mean ± S. E. M. The small error bars indicate tight regulation of Pax7^+^ cell numbers. Number of embryos scored is indicated within the columns. **E:** Co-localization of Pax7 and EdU in MPCs of 4 dpf larva both at VM (white arrowheads) and CP (yellow arrowheads). Vertical myoseptum (VM), central portion (CP), horizontal myoseptum (HM), dorsal edge (DE). Bars 50 μm.

To understand how Pax7^+^ cells arise in the deep somite, the locations of Pax7^+^ cells were characterized at successive stages. In 3 dpf larvae, about half the Pax7^+^ cells were at the superficial VM, mostly oriented with their long axes parallel to the VM (Fig. 2A-C). Some Pax7^+^ cells were in the superficial CP and DE regions, and a few were located deep within the somite at the HM (Fig. 2A,C,D). By 6 dpf, in contrast, Pax7^+^ cell numbers had risen significantly in the deep CP, the central myotome, where the cells were aligned between muscle fibres (Figs 1F,2D; p < 0.001) and within the deep VM (Figs 1F,2C, p < 0.001). The long axes of some Pax7^+^ nuclei in deep VM were not parallel to the VM, but pointed into the somite, suggesting these cells may move between the VM and CP. Although the number of Pax7^+^ cells increased significantly in the deep VM and CP by 6 dpf, no change was detected in superficial VM or CP (Fig. 2C,D; p = 0.302 and 0.942, respectively). Further, the number of Pax7^+^ cells in HM and DE was unchanged from 3 to 6 dpf (Fig. S3). These data show that the number of Pax7^+^ cells increases at specific somitic locations during larval muscle growth and their orientation is suggestive of an inward movement.

The zebrafish DM contains proliferating Pax7^+^ cells (Hammond et al., 2007; Stellabotte et al., 2007), which could act as a source of cells entering the deep myotome. Analysis of Pax7^+^ cell proliferation with a 3 h EdU labelling pulse showed that around 20% Pax7^+^ cells are in S-phase in most somitic regions at 3 and 4 dpf, when cells are beginning to enter the deep myotome (Figs 2E and S4). Proliferation therefore contributes to the increase in Pax7^+^ cells, but the increase in Pax7^+^ cells in the deep myotome does not arise from localised proliferation.

### Pax7^+^ cells migrate into the somite and proliferate

In amniotes, Pax7^+^ cells of the dermomyotome have been shown to enter the myotome when the central dermomyotome disperses (Ben-Yair and Kalcheim, 2005; Gros et al., 2005; Kassar-Duchossoy et al., 2005). To visualize the dynamics of Pax7^+^ cells in live growing fish, a *pax7a* reporter transgene *Tg(Pax7a: EGFP)^MPIEB^* was bred onto a *pfeffer^tm236b^* background to remove xanthophores, which would otherwise express Pax7 (Alsheimer, 2012; Mahalwar et al., 2014; Minchin and Hughes, 2008; Odenthal et al., 1996). *Pax7a:* GFP^+^ cells and Pax7^+^ nuclei were largely co-localised at 3.25 dpf (Fig. S5), and GFP^+^ cells form muscle fibres, consistent with our findings in the regeneration context (Knappe et al., 2015; Pipalia et al., 2016).

Time-lapse confocal analysis of live *pax7a:GFP;pfe^tm236b/tm236b^* larvae showed many *pax7a:GFP*^+^ cells oriented parallel to the VM at 3.5 dpf (Fig. 3A). Occasional VM cells extended into the myotome parallel to adjacent fast fibres, migrated into the deep myotome and some subsequently divided (Fig. 3A). Tracking of all GFP^+^ cells in one epaxial myotome from 3.5 to 4 dpf revealed that while a few cells moved medially up to 20% of somite width, most moved little (Fig. 3B). Nevertheless, rather few were carried away from the midline, despite the thickening of the myotome (compare Figs 1C and 3B). GFP^+^ cells were observed to enter the myotome from VM and DE (Fig. 3C, Movie S1), but not directly from the central region of the DM. Few GFP^+^ cells were observed deep in the somite at 3 dpf, but such cells were readily detected from 4 dpf onwards (Fig. 3). We conclude that proliferation and migration of cells from the VM and DE contribute to the rise in Pax7^+^ cells within the deep CP. As the number of Pax7^+^ cells in the superficial somite and VM is undiminished over the period studied, we suggest that proliferation of Pax7^+^ cells is sufficient to replenish the loss of cells from these pools following their migration into the deep CP.

**Fig. 3.**
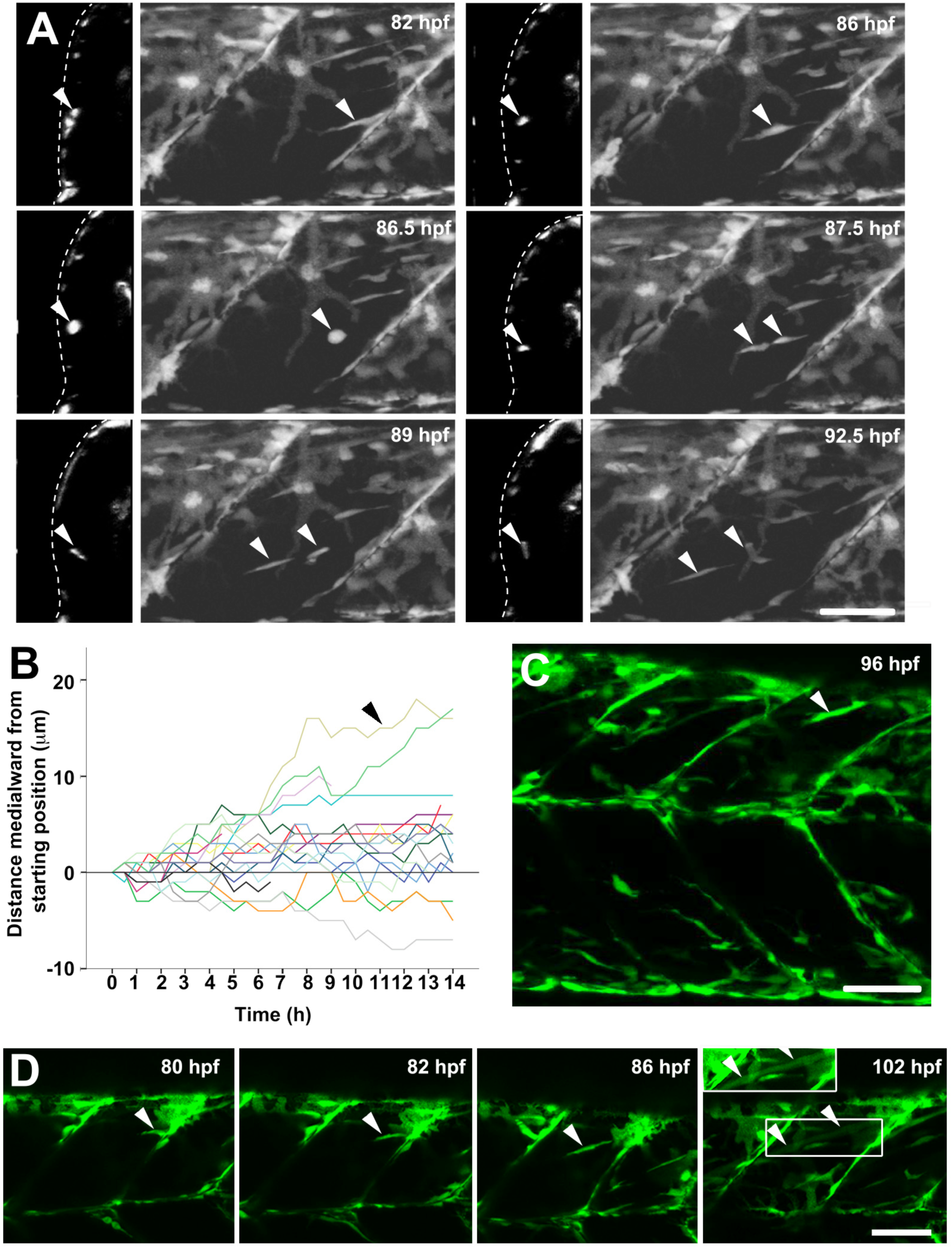
***pax7a:GFP^+^* cells migrate into deep central myotome and differentiate**. Confocal maximum intensity *Z* projections and orthogonal *YZ* views of live *pax7a:GFP*;*pfe/pfe* larvae in lateral view. **A.** Timelapse of 3.5-4 dpf larva taken every 30 min for 14 h showing migration of a cell from the posterior vertical myoseptum into the deep myotome (arrowheads). Note rounding up, division and separation of daughters. **B.** Analysis of distance moved towards midline in *Z* plane for each cell in the epaxial somite from the timelapse shown in A. Note that total movement of cells is often greater, as migration in the anteroposterior and dorsoventral planes (*XY*) is not shown. Each cell was measured relative to its starting position. Arrowhead indicates the cell highlighted in A. **C.** A cell extending from the dorsal edge into the deep myotome (arrowhead). **D.** Timelapse of myotube formation from a cell entering the myotome from the posterior VM (arrowheads). Between 86 and 102 hpf the GFP became diluted in the extended fibre (box is shown with contrast enhancement in inset above). Bars 50 μm.

### Pax7^+^ cells make muscle

Pax7^+^ MPCs give rise to muscle fibres in amniotes (Kassar-Duchossoy et al., 2005; Lepper and Fan, 2010). Neither in fish nor amniotes, however, has Pax7 mRNA or protein been reported in fibres themselves. Time-lapse analysis of *pax7a:GFP* fish showed that Pax7^+^ cells occasionally formed fibres with weak GFP (Fig. 3D). More sensitive immunodetection revealed elongated GFP^+^ fibre-like structures containing sarcomeric MyHC at 4 dpf (Fig. 4A). Thus, perdurant GFP proved that Pax7^+^ cells contribute to muscle growth.

**Fig. 4.**
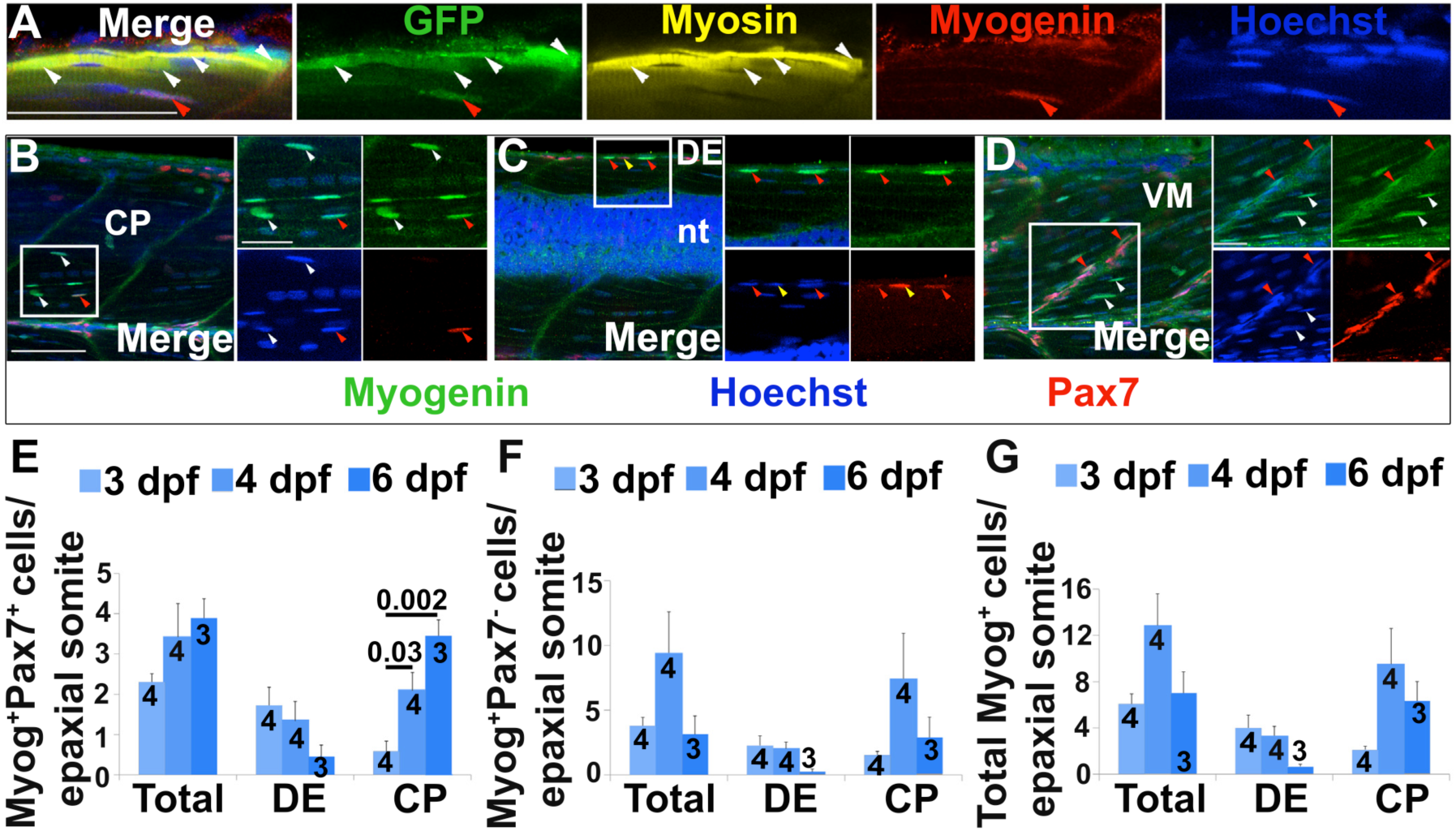
**Pax7^+^ cells differentiate in specific somite regions.** Single confocal planes of 4 dpf wholemount immunofluorescence in lateral view. Scale bars 50 μm. **A.** *Pax7a:GFP*;*pfe/pfe* larva showing GFP in a MyHC^+^ muscle fibre (white arrowheads). A deep GFP^+^MyHC^−^ cell co-labels with Myogenin (red arrowheads). **B-D.** Pax7 and Myogenin in epaxial somite of wt larva showing Pax7^+^Myog^+^ cells (red arrowheads) in CP (B), DE (C) and VM (D), Pax7^+^Myog^−^ cells (yellow arrowheads) in DE (C) and Pax7^−^Myog^+^ cells (white arrowheads) in CP (B, D). Note the reduced Pax7 and Myog signal in Pax7^+^Myog^+^ cells. **E-G.** Time course and location of Myog^+^Pax7^+^ (E), Myog^+^Pax7^-^ (F) and total Myog^+^ (G) cells.

Some GFP^+^ mononucleate cells in the myotome were found to contain Myogenin (Fig. 4A), a marker of differentiating myoblasts in amniotes and embryonic zebrafish (Hasty et al., 1993; Hinits et al., 2009; Nabeshima et al., 1993). Dual immunodetection of Myog and Pax7 proteins in epaxial somites between 3 and 6 dpf confirmed Pax7^+^Myog^+^ cells in CP, DE and more rarely in VM (Fig. 4B-D, respectively). Similarly, Pax7^−^Myog^+^ cells were predominantly in the CP, occasionally at the DE, rarely at HM and were not observed at VM (Fig. 4B-D and data not shown). Pax7^+^Myog^+^ cells tended to show weaker Pax7 and Myog labelling than in the respective single-positive cells (Fig. 4B-D), suggesting a transition between Pax7^+^Myog^−^ and Pax7^−^Myog^+^ cells. The predominant locations of Myog^+^ cells in CP and DE suggest these are the major regions of myoblast terminal differentiation.

Counting Myog^+^ cells revealed a transition in muscle differentiation between 3 and 6 dpf. At 3 dpf, most Myog^+^ cells were located in the DE. From 4 dpf onwards, most Myog^+^ cells were in the CP (Fig. 4E-G). The total number of Pax7^+^Myog^+^ cells increased between 3 and 6 dpf (p=0.02, Fig. 4E), primarily due to an increase in Pax7^+^Myog^+^ cells in the CP (p=0.002); there was no change at the DE (p=0.08). These results indicate that many of the Pax7^+^ cells that invade CP myotome rapidly differentiate and contribute to myotome growth. We conclude that although terminal differentiation removes Pax7^+^ cells, proliferation and migration is sufficient to replenish the Pax7^+^ cell pool.

### Altered growth in myod^fh261^ mutants

Embryos lacking *myod* function show a 50% reduction in muscle at 1 dpf accompanied by a twofold excess of MPCs. This followed by rapid growth resulting in recovery by 5 dpf (Hammond et al., 2007; Hinits et al., 2011). Recovery did not, however, lead to normal muscle. Analysis of fibre number and size revealed that at 5 dpf *myod^fh261^* mutants have 33% fewer fibres, but these are about 30% larger than those in wt (Fig. 5A-C). Thus, recovery compensated for the reduced fibre number by increased fibre growth.

**Fig. 5.**
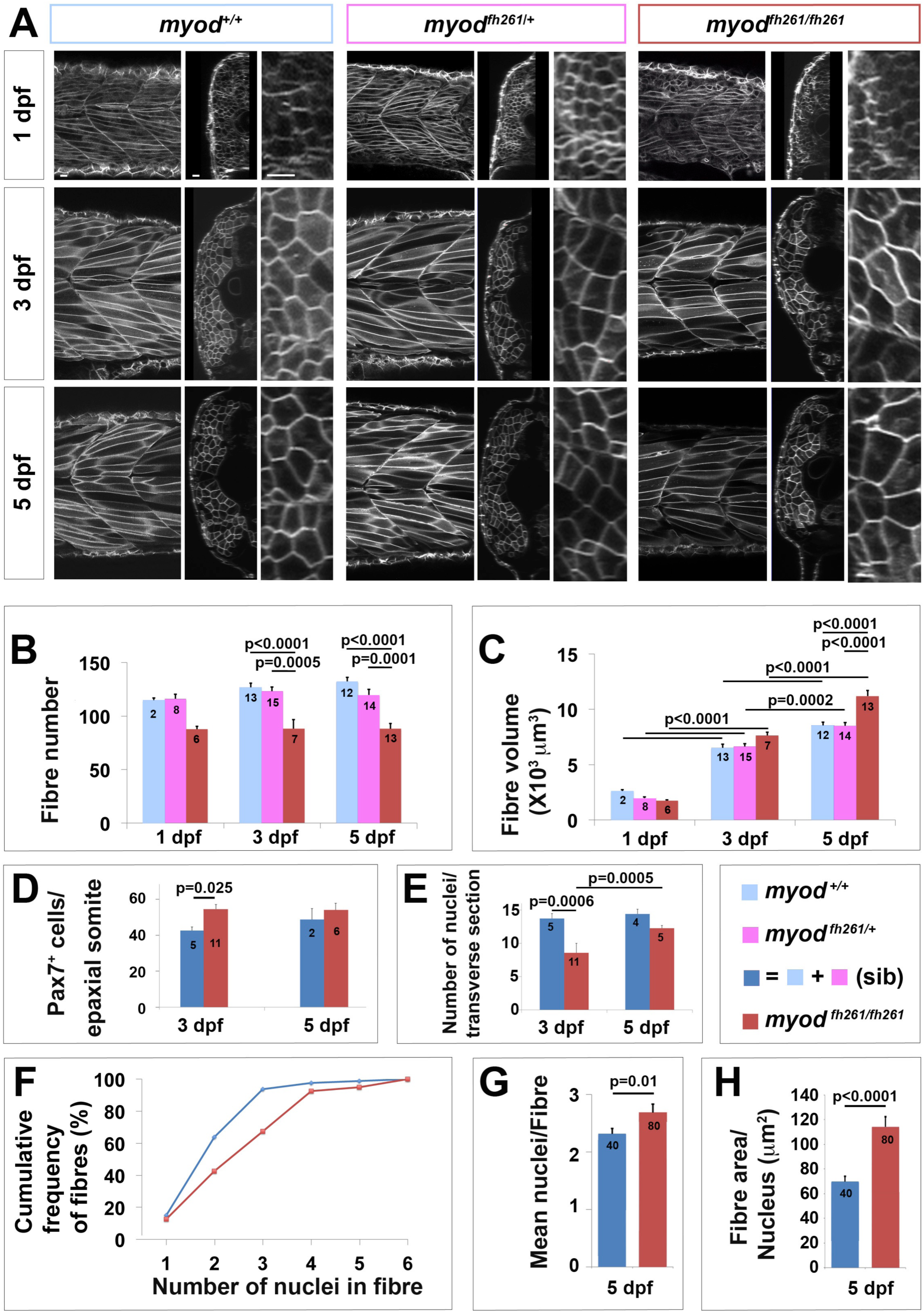
**Lack of *myod* alters somite growth. A.** Single confocal planes of *myod^fh261^* mutant and sibling zebrafish expressing plasma membrane-GFP from *Tg(Ola.Actb:Hsa.HRAS-EGFP)^vu119^*. Individual retrospectively genotyped larvae are shown at successive stages in each tryptich in lateral (left), transverse (centre) and magnified transverse (right) views. Bars 10 μm. **B.** Fibre number per somite in each stage and genotype as indicated in key. **C.** Mean fibre volume = myotome volume/fibre number. **D, E.** Larvae from *myod^fh261^* heterozygote incross were stained at 3 and 5 dpf for Pax7, MyHC and DNA and analysed by confocal microscopy. Pax7^+^ cell number/epaxial somite (D) and number of nuclear profiles within the myotome/transverse optical section of epaxial somite 17 (E) were scored from fish genotyped by loss of head myogenesis. **F-H.** Analysis of nuclear number in fast fibres of *myod^fh261^* mutant and sibling *Tg(Ola.Actb:Hsa.HRAS-EGFP)^vu119^* injected with RNA encoding H2B-mCherry to permit counting in live larvae. Fibres analysed (number on columns) are shown in Fig. S6. A cumulative frequency plot (F) reveals the larger number of nuclei in fibres of mutants, which is reflected in an average of 16% increase in nuclear number (G) accompanied by a 63% increase in fibre cross-sectional area per nucleus (H), reflecting an 80% increase in fibre size. Differences tested by ANOVA with Tukey post-hoc (B, C), Kruskall-Wallis (F, G) and t-test (E, H).

Analysis of fibre number and size during muscle recovery showed significant defects in *myod^fh261^* mutants. In wt embryos, fibre numbers increased by about 15% from 1-5 dpf. No increase was observed in *myod^fh261^* mutants (Fig. 5B). In contrast, fibre size was comparable in wt and *myod^fh261^* mutants at 1 dpf, but fibre size increased faster in mutants so that, by 5 dpf, fibres were larger, thereby compensating for the reduction in fibre number (Fig. 5C).

*Myod^fh261^* mutants have an increased number of Pax3/7^+^ cells at 1 dpf, paralleling the reduction in muscle differentiation (Hinits et al., 2011). The number of Pax7^+^ cells remains elevated at 3 dpf, but returns almost to normal by 5 dpf, accompanied by a rise in the number of myonuclei in the myotome (Fig. 5D,E).

Do *myod^fh261^* mutants recover by fusion of the excess myoblasts into the pre-existing fibres? Both the maximal number and the mean number of nuclei in single fast muscle fibres of *myod^fh261^* mutants was higher than in siblings (Figs 5F,G and S6), suggesting that the excess myoblasts contribute to the growth of existing myotomal fibres. However, when the fibre size per nucleus was calculated (by dividing the cross-sectional area of each fibre by its nuclear number), *myod^fh261^* mutants had a clear increase in effective nuclear domain size (Figs 5H and S6). Thus, fusion of excess Pax7^+^ DM cells into pre-existing fibres during recovery of *myod^fh261^* mutants accompanies hypertrophy – an increase in fibre volume per nucleus.

### Excess Pax7^+^ cells in the deep myotome of 3 day myod^fh261^ mutants

To understand the contribution of DM cells to the recovery of *myod^fh261^* mutants the number and location of Pax7^+^ and Myog^+^ cells were determined (Fig. 6). In 3 dpf *myod^fh261^* mutants there were approximately 25% more Pax7^+^ cells than in wt (p=0.025, Fig. 6A-C). Strikingly, the extra Pax7^+^ cells in mutants at 3 dpf were mostly located in the deep CP myotome (Fig. 6B-E. This reveals an earlier presence of Pax7^+^ cells in the deep myotome of *myod^fh261^* mutants than in siblings and wt fish, which accumulate such cells from 4 dpf (Fig. 3).

**Fig. 6.**
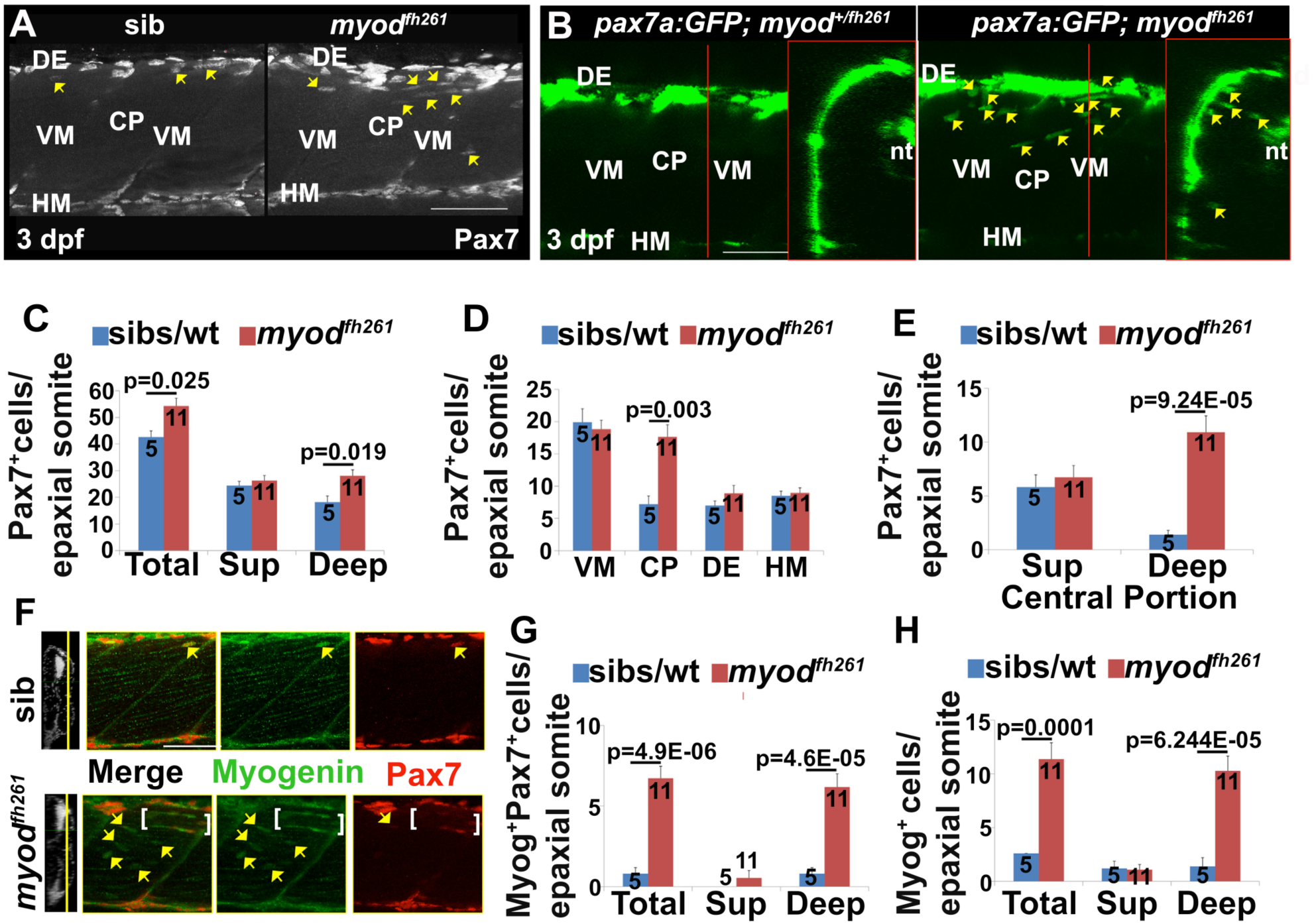
**Premature ingression of Pax7^+^ myogenic precursors in *myod^fh261^* mutants**. Wholemount larvae from a *myod^fh261/+^* incross stained for Pax7 (A, C-H) or *pax7a:GFP;myod^fh261/+^* incross imaged live (B) at 3 dpf and shown in confocal short stacks in lateral view. **A.** Pax7 antibody stained nuclei (arrows) within the deep CP myotome. **B.** Pax7:GFP^+^ cells (arrows) in lateral and transversal views. **C-H.** Comparison of Pax7^+^ (C-E), Myog^+^Pax7^+^ (G) and Myog^+^ (H) cell numbers in the epaxial half of som16-18 between the number of *myod^fh261^* mutants and their siblings indicated within columns. The extra Pax7^+^ cells in *myod^fh261^* mutants (C) were specifically located in deep CP (D, E). Lateral planes at deep locations indicated by yellow line on transversal sections of wholemount larvae stained for Pax7, Myogenin and Hoechst (F) reveal increased numbers of Myog^+^ and Pax7^+^Myog^+^ in the deep CP (arrows). Note the alignment of some Myog^+^ nuclei (brackets). VM: vertical myosepta, CP: central portion, DE: dorsal edge, HM: horizontal myoseptum. Bars 50 μm.

The extra Pax7^+^ cells in 3 dpf *myod^fh261^* mutants appear to be undergoing differentiation. More Myog^+^ cells were observed in the deep CP of *myod^fh261^* mutants at 3 dpf, compared with their siblings (Fig. 6F-H). Moreover, these increases were observed specifically in the deep central myotome (Fig. 6G,H), and not at other locations of the myotome. Comparing *myod^fh261^* mutants and siblings, a similar fraction of Pax7^+^ cells were also Myog^+^ and the ratio of Pax7^+^Myog^+^ to Pax7^−^Myog^+^ cells was unaltered between genotypes. These findings suggest that, once in the deep central myotome, *myod^fh261^* mutant Pax7^+^ cells progress to terminal differentiation in the normal manner. At 5 dpf, no significant difference in either Pax7^+^ or Myog^+^ cell numbers persisted (Fig. S7A-C). These data argue that the premature appearance of Pax7^+^ cells in the deep central myotome does not reflect a failure of differentiation in *myod^fh261^* mutant, but rather an adaptive process contributing to increase in nuclei/fibre and muscle mass recovery.

If the increase in MPCs in the deep central myotome reflects an adaptive process, it could arise either from increased migration or proliferation of Pax7^+^ cells. To examine this issue, *myod^fh261^* was crossed onto the *pax7a:GFP* transgene to permit tracking of cell dynamics. Profiles suggesting migration of cells from the vertical myosepta into the deep myotome were more common at 3 dpf in *myod^fh261^* mutants than in siblings (Fig. 6B and data not shown). Moreover, EdU labelling showed that cell proliferation was similar in the deep central myotome in mutants and siblings (Fig. S7D-F). We conclude that there is an increased migration of Pax7^+^ cells into the deep myotome of 3 dpf *myod^fh261^* mutants.

### Blockade of GSK3 reduces accumulation of Pax7 cells in the myotome

A small molecule screen for pathways that affect Pax7^+^ cell behaviour revealed that GSK3 signalling may regulate migration (Fig. 7A,B). GSK3 is downstream of various signalling pathways, including Wnt/β-catenin and Insulin, both of which can influence muscle growth (Bentzinger et al., 2014; Fernandez et al., 2002; Jones et al., 2015; Musaro et al., 2001; Tee et al., 2009). Treatment of wt larvae at 3 dpf with the GSK3 antagonist (2′Z,3′E)-6- bromoindirubin-3′-oxime (BIO) for 24 hours reduced the number of Pax7^+^ cells in the deep CP myotome compared with vehicle (Fig. 7A). Quantification revealed that the number of GFP^+^ cells was significantly decreased in the deep CP of BIO-treated larvae, but relatively unaffected elsewhere (Fig. 7B). The numbers of differentiating Pax7^+^ cells and Myog^+^ cells were also significantly reduced in both superficial and deep CP (Fig. S8). Importantly, BIO also blocked the premature entry of Pax7^+^ cells into the myotome in *myod^fh261^* mutants (Fig. 7C). Thus, BIO prevents migration and/or accumulation of Pax7a^+^ cells in the deep CP and alters terminal differentiation.

**Fig. 7.**
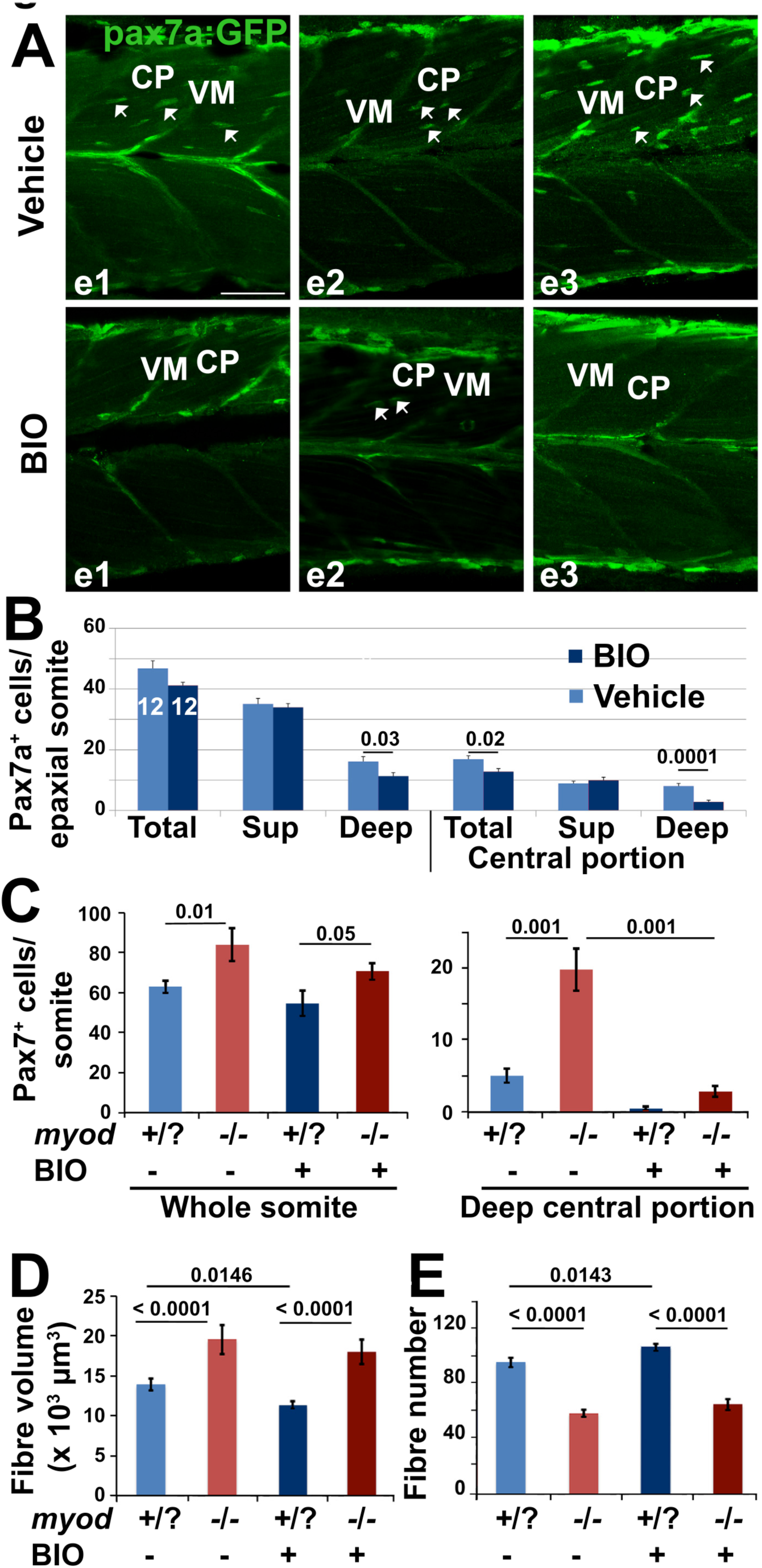
**Blockade of ingression of Pax7a^+^ cells by BIO. A.** The *pax7a:GFP* BAC transgene was bred onto a *pfe/pfe* background to diminish xanthophores and larvae at 3.25 dpf treated with BIO or vehicle were stained in wholemount at 4.25 dpf for GFP, Myog and DNA. Confocal images are maximum intensity projections of short stacks in the deep myotome in lateral view with dorsal up and anterior to left, showing the decline of GFP^+^ cells (arrows) between fibres in the deep central myotome of three individual embryos in each condition. VM vertical borders, CP central portion. Bar 50 μm. **B.** Number of pax7a:GFP^+^ cells in epaxial region of somites 16-18 of 12 BIO-treated embryos compared with 12 controls. Significantly fewer Pax7a^+^ cells were present deep within the somite, which was accounted for by highly significant loss from the deep CP. Sup = superficial. **C.** Larvae from an incross of *pax7a:GFP;myod^fh261/+^* were sorted for GFP, treated at 2.25 dpf with BIO or vehicle, stained at 3.25 dpf for GFP, Pax7 and myosin and genotyped by head muscle. Total (left) and deep CP (right) Pax7^+^ cells were counted from confocal stacks of five larvae in each condition and differences tested by ANOVA with Bonferroni multiple comparison test. **D, E.** Larvae from an incross of *myod^fh261/+^*;*Tg(Ola.Actb:Hsa.HRAS-EGFP)^vu119^* were sorted for GFP, treated with BIO or vehicle from 2-3 dpf and analysed live at 4 dpf by confocal microscopy for mean fibre volume (D) and fibre number (E). Each larva was retrospectively sequence genotyped.

### BIO fails to reduce compensatory muscle fibre growth

As GSK3 inhibition led to fewer Pax7^+^ cells migrating into the deep myotome, it was possible to investigate the importance of this migration in recovery of the muscle in *myod^fh261^* mutants. When *myod^fh261^* mutant embryos were treated with BIO at 2 dpf, thus blocking the premature ingression of Pax7^+^ cells to the deep CP, the increase in muscle fibre volume triggered by the loss of Myod still occurred (Fig. 7D). Interestingly, BIO caused a slight rise in fibre number in wt siblings, but in *myod^fh261^* mutants there was no significant change in fibre number (Fig. 7E). These data indicate that hypertrophy of fibres by increase in volume per nucleus provides the robust recovery of muscle in *myod^fh261^* mutants and migration of Pax7^+^ cell from DM is not required for this recovery.

## Discussion

The present work demonstrates that Pax7^+^ MPCs contribute to larval muscle growth and contains four major findings relevant to myogenesis. First, we show that Myod function is required for formation of the correct number of fast muscle fibres. Second, we describe regional variation in the behaviour of MPCs that likely reflect local control of their proliferation and subsequent differentiation during development. Third, we demonstrate a role for GSK3 signalling in regulating MPC migration, and possibly differentiation, within the early somite. Fourth, we find that muscle stem and precursor cell populations in the early somite are precisely regulated in response to perturbations so as to return the system rapidly to near-normal status, yielding developmental robustness. Our observations suggest that much of the control of myogenesis, even in a simple system, remains unexplained and introduce the zebrafish myotome as a single cell-resolution vertebrate model for the quantitative study of tissue growth and maintenance.

### Regulation of fibre number

Fish lacking Myod fail to form one third of the normal number of somitic fast muscle fibres. Initially, *myod^fh261^* mutants also have an excess of MPCs, suggesting that reduced differentiation explains the reduced fibre number. Although mutants recover muscle volume through fibre hypertrophy, fast fibre number does not recover. Despite the early reduction in total myonuclei in *myod^fh261^* mutants, they show partial recovery by 5 dpf. Thus, the recovery of myotome volume previously reported in *myod^fh261^* mutants (Hinits et al., 2011) involves both hypertrophy and elevated fusion of MPCs, leading to more nuclei in the enlarged fast fibres. Although some excess MPCs differentiate, they fail to make new fibres. This argues for a critical window in myotomal fast muscle development during which the first fibre cohort is initiated and the number of its fibres determined.

In fish, a prolonged initial period of polarized hyperplasia, in which new fibres are made at the dorsal and ventral edges of the myotome, leads to myotome growth. This is followed, in many species, by mosaic hyperplasia, in which new fibres are formed between existing fibres in a manner similar to amniote secondary fibre formation. Environmental changes that alter fish behaviour also affect fibre formation (Johnston et al., 2009; Johnston et al., 2003; Macqueen et al., 2008), but it is unclear whether either polarized or mosaic hyperplasia is nerve-dependent (Johnston et al., 2011). One study that examined the relationship of initial fibre number and final muscle mass revealed that final fibre number is under genetic selection, but did not determine at what point in development the regulatory genes act (Johnston et al., 2004). We observed a failure of polarized hyperplasia in *myod^fh261^* mutant larvae. Future studies in larval zebrafish, where the limited MPC and fibre numbers make quantification practicable, may provide deeper insight into the control of fibre number.

### Regional variations in division, migration and differentiation of MPCs

MPCs in wt fish behave distinctly depending on their somitic location. Each epaxial somite has about 40 Pax7^+^ MPCs at 3 dpf, a number that correlates well with the number of nuclei present in the somite that are not part of fibres (S. Hughes, unpublished observation), but one that is higher than previously reported for whole somites (Seger et al., 2011) Published values for Pax7^+^ cells per somite vary widely even at 24 hpf (Feng et al., 2006; Hammond et al., 2007; Hollway et al., 2007; Seger et al., 2011); our light fixation regime and long antibody incubations may explain the difference. The regional variations in MPC proliferation (reduced at HM), migration (primarily from VM into myotome) and differentiation (highest in DE and deep myotome) that we describe show that MPC dynamics are balanced and locally controlled. Little is known of local signals in zebrafish larvae, although Fgf, Hh and Wnt signalling have been implicated in controlling DM behaviour at earlier stages (Feng et al., 2006; Hammond et al., 2007; Lewis et al., 1999; Tee et al., 2009). To understand muscle growth, it will be essential to elucidate how local signals affect myogenesis in discreet somitic regions.

### Role of GSK3 signalling in myogenesis

Our small molecule screen revealed that inhibition of GSK3 using the ATP-competitor BIO prevents the normal and induced migration of a subset of MPCs into the myotome. GSK3 regulates several signalling pathways, including Wnt/βcat. In vitro, BIO has been shown to maintain human embryonic stem cells in the pluripotent state and prevent their epithelial-mesenchymal transition by mimicking the action of the Wnt/ßcat pathway (Sato et al., 2004; ten Berge et al., 2011; Ullmann et al., 2008). Wnt signalling through GSK3ß can regulate DM behaviour in amniotes and Wnt signals also control muscle fibre patterning in the amniote somite (Gros et al., 2009; Hutcheson et al., 2009; Linker et al., 2005). Once myoblasts have formed, β-catenin-dependent Wnt signalling has been suggested to promote their differentiation (Brack et al., 2008; Jones et al., 2015), whereas non-canonical Wnt signalling may expand MPCs and promote their motility (Bentzinger et al., 2014; Le Grand et al., 2009). We observed a small but significant increase in fibre number in response to BIO, suggesting that some MPCs have differentiated to form new fibres. Thus, as in amniotes, GSK3ß activity may play different roles in successive stages of myogenesis (Lien and Fuchs, 2014; Murphy et al., 2014).

In zebrafish, gain of function Wnt/β-catenin signalling caused by *axin1*/*apc1* double mutation has been shown to increase Pax3/7 cell proliferation, but without increase in Pax3/7 cell number (Tee et al., 2009). Congruently, we observe that BIO prevents entry of Pax7a^+^ cells into the myotome, also without a significant increase in dermomyotome cell number. These findings may be explained by the increased dermomyotomal apoptosis observed by Tee et al. (2009). Tee et al. also report increased apparent thickness of fibres in *axin1*/*apc1* mutants and LiCl-treated embryos. We observe no such effect with BIO applied at a somewhat later stage, or with LiCl treatment (J. Groves and S. M. Hughes, unpublished data). On the contrary, if anything, we observe a slight reduction in myotome volume by treatment with BIO alone, but no effect on the recovery of the myotome in *myod* mutants. Currently, we favour the simple interpretation that nuclear increase in *myod^fh261^* mutant fast fibres arises from increased fusion of MPCs. A definitive test of this hypothesis will require ablation of the various MPC populations.

β-catenin was shown to be required for muscle hypertrophy fibre-autonomously following muscle overload in rodents (Armstrong and Esser, 2005; Armstrong et al., 2006). Our data show that BIO, which is expected to activate β-catenin signalling throughout the fish, does not alter developmental fibre growth. Thus, β-catenin signalling alone may be insufficient to promote fibre hypertrophy.

### Robustness in Myogenesis

The finding that Myod function limits muscle fibre size in larval zebrafish parallels data in mouse showing that Myogenin and Mrf4 can prevent atrophy or induce hypertrophy, respectively (Moresi et al., 2010; Moretti et al., 2016). This emerging theme suggests that a function of the duplicated vertebrate MRF genes within fibres is robust regulation of muscle size. Although Myod is well known to promote MPC terminal differentiation (Kablar et al., 1997; Yablonka-Reuveni et al., 1999), our data are inconsistent with a simple delay in myogenesis in *myod* mutants. Fibre number is reduced, but does not recover when excess MPCs subsequently differentiate, suggesting that a critical window for fibre formation has been missed. Moreover, ongoing new fibre formation is inhibited, raising the possibility that MPCs in specialised regions or with specific characteristics may be particularly vulnerable to lack of Myod function. Instead of a persistent defect, however, overall muscle size regulates to the normal value. This homeostasis is achieved by changing behaviour of both MPCs, through altered migration, and fibres themselves, by increasing volume per nucleus. The recent discovery of MPC diversity in the early zebrafish somite (Gurevich et al., 2016; Pipalia et al., 2016) raises the possibility that distinct MPC populations may preferentially contribute to hypertrophy in *myod* mutants. Such robustness may explain how stochastic variation in the low MPC numbers in each somite does not lead to maladaptive variation in muscle mass along the body axis or between left and right sides.

In summary, we examined the role and regulation of Pax7^+^ MPCs in larval muscle growth. We observed tight regional control of MPC numbers, distribution and behaviour within the somite and myotome. Perturbations that alter muscle size and MPC number were rapidly corrected, suggesting the existence of a homeostatic mechanism that senses muscle size and ensures robust development in the face of environmental and genetic insults.

## Materials and Methods

### Zebrafish lines and maintenance

Genetically-altered *Danio rerio* (listed in Table S1) on a primarily AB background were reared at King’s College London on a 14/10hr light/dark cycle at 28.5°C (Westerfield, 2000). BIO (0.5 μM) or DMSO vehicle were added to fish water.

### Immunodetection and S-phase labelling

Fibre sizes on photomicrographs of cryosections either unstained or after immunoperoxidase detection of MyHC were quantified with OpenLab (Improvision). For wholemounts, larval pigmentation was suppressed with 0.003% 1-phenyl-2-thiourea (Sigma) added at 12 hpf. Larvae were fixed with 2% PFA for 25 minutes, washed with PBTx (PBS, 0.5% or 1% (4 dpf+) Triton-X100) and incubated in primary antibody (see Table S2) for 3-5 days at 4°C on a rotary shaker, washed repeatedly in PBTx, incubated with subclass specific Alexa-conjugated secondary antibodies (Molecular Probes) overnight, repeatedly washed with PBTx prior to incubation with 1 μM Hoechst 33342 for 2 hours at room temperature. Larvae were washed and mounted under a cover slip with Citifluor AF1 for imaging. Larvae were S-phase labelled by exposure to 1 mg/ml EdU in 10% PBS:90% system water for 3 hours, immediately fixed with 2% PFA and EdU detected with a Click-iT kit (Invitrogen C10084).

### Imaging and quantification

All images of fish are oriented dorsal up in either transverse or lateral anterior to left view. Larvae were anaesthetized with MS222, mounted in 1% low melting agarose and viewed laterally by a Zeiss 20x/1.0 NA dipping objective on an LSM Exciter confocal microscope with ZEN (2009+2012) software or a Nikon D-Eclipse C1 microscope with 40x/0.8 NA water dipping objective and EZ-C1 3.70 software. In Fig. 3A, Z-stacks were acquired in 1 μm steps from epidermis to neural tube, processed with Fiji, drift adjusted with ‘Correct 3D drift’ and single cells tracked manually with ‘MtrackJ’. To account for drift and growth, a reference point on the epidermis was also tracked and the respective *Z*-value subtracted from that of individual *pax7a:GFP+* cells at each of the 28 time-points to obtain a depth measurement relative to epidermis. Absolute movement in *Z* for each cell in Fig. 3B was calculated by subtracting the position at 82 hpf from that at each subsequent time-point. Somite volume and fibre number were measured from confocal stacks of somite 17 in *Tg(Ola.Actb:Hsa.HRAS-EGFP)^vu119^* larvae as described (Hinits et al., 2011). Mean fibre number was calculated from three optical sections after correcting for double counting at vertical myosepta (VM) using Fibre number = Total fibre profiles – (Profiles touching VM)/2. Mean fibre volume = Myotome volume/Mean fibre number.

Fixed fish were imaged using the 10x/0.3 air or 40x/1.1 water immersion objectives. Three to nine somites around the anal vent were imaged from lateral using the tile scan *Z*-stack function. Short stack maximum intensity projections, specific slices or cross-sectional views were exported as tiffs. Nuclear number was determined from three equi-spaced transverse images from somite 17 of each embryo. Cells were counted in original ZEN stacks and allocated to regions (Fig. S1) in confocal stacks of epaxial somites of wholemount fish by scanning through in the XZ direction while toggling channels. Xanthophores were excluded from Pax7 counts based on nuclear shape, location and intensity (Hammond et al., 2007).

### Statistics

Statistics were analysed with Microsoft Excel and AnalySoft Statplus, Graphpad Prism 6 or SPSS on the number of samples indicated. F-test was used to determine equivalence of variance and the appropriate Student’s t-test or ANOVA with Scheffé post hoc test applied unless otherwise stated. All graphs show mean and standard error of the mean. Numbers on columns represent number of fish scored.

## Acknowledgements

We thank Jana Koth, Yaniv Hinits and members of the Hughes lab for advice, C. Houart, P. W. Ingham, S. Alsheimer and C. Nüsslien-Volhard for fish lines, Bruno Correia da Silva and his staff for care of the fish.

## Competing Interests

The authors declare no competing interests.

## Author Contributions

Experiments were performed by SDR (Figs 1C,E,F;2;3C,D;4;6A,C-H;7A,B;S1-S5;S7;S8), VCW (Figs 5;7D,E;S6), TGP (Figs 5F-H;6;7C-E;S7), CLH (Fig. 1A,B), KL (Fig. 1D), SK (Fig. 3A,B) and RDK (Fig. 7A,B). RDK proposed and initiated the BIO experiments. SMH conceived the project, provided advice and wrote the manuscript with input from all authors.

## Funding

SMH is a Medical Research Council (MRC) Scientist with Programme Grant G1001029 and MR/N021231/1 support. CLH had a MRC PhD studentship. RDK was funded by the Biotechnology and Biological Sciences Research Council (BB/I025883/1) and Wellcome Trust (101529/Z/13/Z).

## References

Abmayr, S. M. and Pavlath, G. K. (2012). Myoblast fusion: lessons from flies and mice. Development 139, 641–656.

Alsheimer, S. (2012). On teleost muscle stem cells and the vertical myoseptum as their niche. pp. 249. Tübingen, Germany: Universität Tübingen.

Armstrong, D. D. and Esser, K. A. (2005). Wnt/beta-catenin signaling activates growth-control genes during overload-induced skeletal muscle hypertrophy. Am J Physiol Cell Physiol 289, C853–859.

Armstrong, D. D., Wong, V. L. and Esser, K. A. (2006). Expression of beta-catenin is necessary for physiological growth of adult skeletal muscle. Am J Physiol Cell Physiol 291, C185–188.

Barresi, M. J., D’Angelo, J. A., Hernandez, L. P. and Devoto, S. H. (2001). Distinct mechanisms regulate slow-muscle development. Current Biol 11, 1432–1438

Ben-Yair, R. and Kalcheim, C. (2005). Lineage analysis of the avian dermomyotome sheet reveals the existence of single cells with both dermal and muscle progenitor fates. Development 132, 689–701.

Bentzinger, C. F., von Maltzahn, J., Dumont, N. A., Stark, D. A., Wang, Y. X., Nhan, K., Frenette, J., Cornelison, D. D. and Rudnicki, M. A. (2014). Wnt7a stimulates myogenic stem cell motility and engraftment resulting in improved muscle strength. J Cell Biol 205, 97–111.

Bentzinger, C. F., Wang, Y. X. and Rudnicki, M. A. (2012). Building muscle: molecular regulation of myogenesis. Cold Spring Harb Perspect Biol 4, a008342.

Brack, A. S., Conboy, I. M., Conboy, M. J., Shen, J. and Rando, T. A. (2008). A temporal switch from notch to Wnt signaling in muscle stem cells is necessary for normal adult myogenesis. Cell stem cell 2, 50–59.

Buckingham, M. and Rigby, P. W. (2014). Gene regulatory networks and transcriptional mechanisms that control myogenesis. Dev Cell 28, 225–238.

Bunger, L., Navajas, E. A., Stevenson, L., Lambe, N. R., Maltin, C. A., Simm, G., Fisher, A. V. and Chang, K. C. (2009). Muscle fibre characteristics of two contrasting sheep breeds: Scottish Blackface and Texel. Meat Sci 81, 372–381.

Cavalier-Smith, T. (2005). Economy, speed and size matter: evolutionary forces driving nuclear genome miniaturization and expansion. Ann Bot 95, 147–175.

Devoto, S. H., Stoiber, W., Hammond, C. L., Steinbacher, P., Haslett, J. R., Barresi, M. J., Patterson, S. E., Adiarte, E. G. and Hughes, S. M. (2006). Generality of vertebrate developmental patterns: evidence for a dermomyotome in fish. Evol Dev 8, 101–110.

Fankhauser, G. (1945). Maintenance of normal structure in heteroploid salamander larvae, through compensation of changes in cell size by adjustment of cell number and cell shape. J Exp Zool 100, 445–455.

Feng, X., Adiarte, E. G. and Devoto, S. H. (2006). Hedgehog acts directly on the zebrafish dermomyotome to promote myogenic differentiation. Dev Biol 300, 736–746.

Fernandez, A. M., Dupont, J., Farrar, R. P., Lee, S., Stannard, B. and Le Roith, D. (2002). Muscle-specific inactivation of the IGF-I receptor induces compensatory hyperplasia in skeletal muscle. J Clinical Invest 109, 347–355.

Gokhale, R. H. and Shingleton, A. W. (2015). Size control: the developmental physiology of body and organ size regulation. Wiley Interdiscip Rev Dev Biol 4, 335–356.

Gros, J., Manceau, M., Thome, V. and Marcelle, C. (2005). A common somitic origin for embryonic muscle progenitors and satellite cells. Nature 435, 954–958.

Gros, J., Serralbo, O. and Marcelle, C. (2009). WNT11 acts as a directional cue to organize the elongation of early muscle fibres. Nature 457, 589–593.

Groves, J. A., Hammond, C. L. and Hughes, S. M. (2005). Fgf8 drives myogenic progression of a novel lateral fast muscle fibre population in zebrafish. Development 132, 4211–4222.

Gurevich, D. B., Nguyen, P. D., Siegel, A. L., Ehrlich, O. V., Sonntag, C., Phan, J. M., Berger, S., Ratnayake, D., Hersey, L., Berger, J., et al. (2016). Asymmetric division of clonal muscle stem cells coordinates muscle regeneration in vivo. Science 353, aad9969.

Hammond, C. L., Hinits, Y., Osborn, D. P., Minchin, J. E., Tettamanti, G. and Hughes, S. M. (2007). Signals and myogenic regulatory factors restrict pax3 and Pax7 expression to dermomyotome-like tissue in zebrafish. Dev Biol 302, 504–521.

Hasty, P., Bradley, A., Morris, J. H., Edmondson, D. G., Venuti, J. M., Olson, E. N. and Klein, W. H. (1993). Muscle deficiency and neonatal death in mice with targeted mutation in the *myogenin gene*. Nature 364, 501–506.

Hinits, Y., Osborn, D. P. and Hughes, S. M. (2009). Differential requirements for myogenic regulatory factors distinguish medial and lateral somitic, cranial and fin muscle fibre populations. Development 136, 403–414.

Hinits, Y., Williams, V. C., Sweetman, D., Donn, T. M., Ma, T. P., Moens, C. B. and Hughes, S. M. (2011). Defective cranial skeletal development, larval lethality and haploinsufficiency in Myod mutant zebrafish. Dev Biol 358, 102–112.

Hollway, G. E., Bryson-Richardson, R. J., Berger, S., Cole, N. J., Hall, T. E. and Currie, P. D. (2007). Whole-somite rotation generates muscle progenitor cell compartments in the developing zebrafish embryo. Dev Cell 12, 207–219.

Hughes, S. M., Chi, M. M., Lowry, O. H. and Gundersen, K. (1999). Myogenin induces a shift of enzyme activity from glycolytic to oxidative metabolism in muscles of transgenic mice. J Cell Biol 145, 633–642.

Hutcheson, D. A., Zhao, J., Merrell, A., Haldar, M. and Kardon, G. (2009). Embryonic and fetal limb myogenic cells are derived from developmentally distinct progenitors and have different requirements for beta-catenin. Genes Dev 23, 997–1013.

Irvine, K. D. and Harvey, K. F. (2015). Control of Organ Growth by Patterning and Hippo Signaling in Drosophila. Cold Spring Harb Perspect Biol 7, pii: a019224.

Johnston, I. A., Abercromby, M., Vieira, V. L., Sigursteindottir, R. J., Kristjansson, B. K., Sibthorpe, D. and Skulason, S. (2004). Rapid evolution of muscle fibre number in post-glacial populations of Arctic charr Salvelinus alpinus. J Exp Biol 207, 4343–4360.

Johnston, I. A., Bower, N. I. and Macqueen, D. J. (2011). Growth and the regulation of myotomal muscle mass in teleost fish. J Exp Biol 214, 1617–1628.

Johnston, I. A., Lee, H. T., Macqueen, D. J., Paranthaman, K., Kawashima, C., Anwar, A., Kinghorn, J. R. and Dalmay, T. (2009). Embryonic temperature affects muscle fibre recruitment in adult zebrafish: genome-wide changes in gene and microRNA expression associated with the transition from hyperplastic to hypertrophic growth phenotypes. J Exp Biol 212, 1781–1793.

Johnston, I. A., Manthri, S., Smart, A., Campbell, P., Nickell, D. and Alderson, R. (2003). Plasticity of muscle fibre number in seawater stages of Atlantic salmon in response to photoperiod manipulation. J Exp Biol 206, 3425–3435.

Jones, A. E., Price, F. D., Le Grand, F., Soleimani, V. D., Dick, S. A., Megeney, L. A. and Rudnicki, M. A. (2015). Wnt/beta-catenin controls follistatin signalling to regulate satellite cell myogenic potential. Skeletal muscle 5, 14.

Kablar, B., Krastel, K., Ying, C., Asakura, A., Tapscott, S. J. and Rudnicki, M. A. (1997). MyoD and Myf-5 differentially regulate the development of limb versus trunk skeletal muscle. Development 124, 4729–4738.

Kassar-Duchossoy, L., Giacone, E., Gayraud-Morel, B., Jory, A., Gomes, D. and Tajbakhsh, S. (2005). Pax3/Pax7 mark a novel population of primitive myogenic cells during development. Genes Dev 19, 1426–1431.

Knappe, S., Zammit, P. S. and Knight, R. D. (2015). A population of Pax7-expressing muscle progenitor cells show differential responses to muscle injury dependent on developmental stage and injury extent. Front Aging Neurosci 7, 161.

Le Grand, F., Jones, A. E., Seale, V., Scime, A. and Rudnicki, M. A. (2009). Wnt7a activates the planar cell polarity pathway to drive the symmetric expansion of satellite stem cells. Cell stem cell 4, 535–547.

Lepper, C. and Fan, C. M. (2010). Inducible lineage tracing of Pax7-descendant cells reveals embryonic origin of adult satellite cells. Genesis 48, 424–436.

Lewis, K. E., Concordet, J. P. and Ingham, P. W. (1999). Characterisation of a second *patched gene* in the zebrafish *Danio rerio* and the differential response of *patched* genes to Hedgehog signalling. Dev Biol 208, 14–29.

Lien, W. H. and Fuchs, E. (2014). Wnt some lose some: transcriptional governance of stem cells by Wnt/beta-catenin signaling. Genes Dev 28, 1517–1532.

Linker, C., Lesbros, C., Gros, J., Burrus, L. W., Rawls, A. and Marcelle, C. (2005). beta-Catenin-dependent Wnt signalling controls the epithelial organisation of somites through the activation of paraxis. Development 132, 3895–3905.

Macqueen, D. J., Robb, D. H., Olsen, T., Melstveit, L., Paxton, C. G. and Johnston, I. A. (2008). Temperature until the ‘eyed stage’ of embryogenesis programmes the growth trajectory and muscle phenotype of adult Atlantic salmon. Biol Lett 4, 294–298.

Mahalwar, P., Walderich, B., Singh, A. P. and Nüsslein-Volhard, C. (2014). Local reorganization of xanthophores fine-tunes and colors the striped pattern of zebrafish. Science 345, 1362–1364.

Maves, L., Waskiewicz, A. J., Paul, B., Cao, Y., Tyler, A., Moens, C. B. and Tapscott, S. J. (2007). Pbx homeodomain proteins direct Myod activity to promote fast20 muscle differentiation. Development 134, 3371–3382.

Minchin, J. E. and Hughes, S. M. (2008). Sequential actions of Pax3 and Pax7 drive xanthophore development in zebrafish neural crest. Dev Biol 317, 508–522.

Minchin, J. E., Williams, V. C., Hinits, Y., Low, S., Tandon, P., Fan, C. M., Rawls, J. F. and Hughes, S. M. (2013). Oesophageal and sternohyal muscle fibres are novel Pax3-dependent migratory somite derivatives essential for ingestion. Development 140, 2972–2984.

Moresi, V., Williams, A. H., Meadows, E., Flynn, J. M., Potthoff, M. J., McAnally, J., Shelton, J. M., Backs, J., Klein, W. H., Richardson, J. A., et al. (2010). Myogenin and class II HDACs control neurogenic muscle atrophy by inducing E3 ubiquitin ligases. Cell 143, 35–45.

Moretti, I., Ciciliot, S., Dyar, K. A., Abraham, R., Murgia, M., Agatea, L., Akimoto, T., Bicciato, S., Forcato, M., Pierre, P., et al. (2016). MRF4 negatively regulates adult skeletal muscle growth by repressing MEF2 activity. Nature Comms 7, 12397.

Musaro, A., McCullagh, K., Paul, A., Houghton, L., Dobrowolny, G., Molinaro, M., Barton, E. R., Sweeney, H. L. and Rosenthal, N. (2001). Localized Igf-1 transgene expression sustains hypertrophy and regeneration in senescent skeletal muscle. Nature Genet 27, 195–200.

Murphy, M. M., Keefe, A. C., Lawson, J. A., Flygare, S. D., Yandell, M. and Kardon, G. (2014). Transiently active Wnt/beta-catenin signaling is not required but must be silenced for stem cell function during muscle regeneration. Stem Cell Reports 3, 475–488.

Nabeshima, Y., Hanaoka, K., Hayasaka, M., Esumi, E., Li, S., Nonaka, I. and Nabeshima, Y.-i. (1993). *Myogenin* gene disruption results in perinatal lethality owing to severe muscle defect. Nature 364, 532–535.

Neyt, C., Jagla, K., Thisse, C., Thisse, B., Haines, L. and Currie, P. D. (2000). Evolutionary origins of vertebrate appendicular muscle. Nature 408, 82–86.

Odenthal, J., Rossnagel, K., Haffter, P., Kelsh, R. N., Vogelsang, E., Brand, N., Van Eeden, F. J. M., Furutani-Seiki, A., Granato, R., Hammerschmidt, M., et al. (1996). Mutations affecting xanthophore pigmentation in the zebrafish, Danio rerio. Development 123, 391–398.

Ontell, M., Hughes, D. and Bourke, D. (1988). Morphometric analysis of the developing mouse soleus muscle. Am. J. Anat. 181, 279–288.

Ontell, M. and Kozeka, K. (1984). Organogenesis of the mouse extensor digitorum logus muscle: a quantitative study. Am. J. Anat. 171, 149–161.

Otto, S. P. (2007). The evolutionary consequences of polyploidy. Cell 131, 452–462.

Patterson, S. E., Mook, L. B. and Devoto, S. H. (2008). Growth in the larval zebrafish pectoral fin and trunk musculature. Dev Dyn 237, 307–315.

Pipalia, T. G., Koth, J., Roy, S. D., Hammond, C. L., Kawakami, K. and Hughes, S. M. (2016). Cellular dynamics of regeneration reveals role of two distinct Pax7 stem cell populations in larval zebrafish muscle repair. Disease Mod Mech 9, 671–684.

Rawls, A., Morris, J. H., Rudnicki, M., Braun, T., Arnold, H. H., Klein, W. H. and Olson, E. N. (1995). Myogenin's functions do not overlap with those of MyoD or Myf-5 during mouse embryogenesis. Dev Biol 172, 37–50.

Relaix, F. and Zammit, P. S. (2012). Satellite cells are essential for skeletal muscle regeneration: the cell on the edge returns centre stage. Development 139, 2845–2856.

Rudnicki, M. A., Braun, T., Hinuma, S. and Jaenisch, R. (1992). Inactivation of MyoD in mice leads to up-regulation of the myogenic HLH gene Myf-5 and results in apparently normal muscle development. Cell 71, 383–390.

Rudnicki, M. A., Schnegelsberg, P. N., Stead, R. H., Braun, T., Arnold, H. H. and Jaenisch, R. (1993). MyoD or Myf-5 is required for the formation of skeletal muscle. Cell 75, 1351–1359.

Sato, N., Meijer, L., Skaltsounis, L., Greengard, P. and Brivanlou, A. H. (2004). Maintenance of pluripotency in human and mouse embryonic stem cells through activation of Wnt signaling by a pharmacological GSK-3-specific inhibitor. Nature Med 10, 55–63.

Schnapp, E., Pistocchi, A. S., Karampetsou, E., Foglia, E., Lamia, C. L., Cotelli, F. and Cossu, G. (2009). Induced early expression of mrf4 but not myog rescues myogenesis in the myod/myf5 double-morphant zebrafish embryo. J Cell Sci 122, 481–488.

Seger, C., Hargrave, M., Wang, X., Chai, R. J., Elworthy, S. and Ingham, P. W. (2011). Analysis of Pax7 expressing myogenic cells in zebrafish muscle development, injury, and models of disease. Dev Dyn 240, 2440–2451.

Stellabotte, F. and Devoto, S. H. (2007). The teleost dermomyotome. Dev Dyn 236, 2432–2443.

Stellabotte, F., Dobbs-McAuliffe, B., Fernandez, D. A., Feng, X. and Devoto, S. H. (2007). Dynamic somite cell rearrangements lead to distinct waves of myotome growth. Development 134, 1253–1257.

Tajbakhsh, S., Rocancourt, D., Cossu, G. and Buckingham, M. (1997). Redefining the genetic hierarchies controlling skeletal myogenesis: Pax-3 and Myf-5 act upstream of MyoD. Cell 89, 127–138.

Tee, J. M., van Rooijen, C., Boonen, R. and Zivkovic, D. (2009). Regulation of slow and fast muscle myofibrillogenesis by Wnt/beta-catenin and myostatin signaling. PLoS One 4, e5880.

tenBerge, D., Kurek, D., Blauwkamp, T., Koole, W., Maas, A., Eroglu, E., Siu, R. K. and Nusse, R. (2011). Embryonic stem cells require Wnt proteins to prevent differentiation to epiblast stem cells. Nat Cell Biol 13, 1070–1075.

Ullmann, U., Gilles, C., De Rycke, M., Van de Velde, H., Sermon, K. and Liebaers, I. (2008). GSK-3-specific inhibitor-supplemented hESC medium prevents the epithelial-mesenchymal transition process and the up-regulation of matrix metalloproteinases in hESCs cultured in feeder-free conditions. Mol Hum Reprod 14, 169–179.

Venuti, J. M., Morris, J. H., Vivian, J. L., Olson, E. N. and Klein, W. H. (1995). Myogenin is required for late but not early aspects of myogenesis during mouse development. J Cell Biol 128, 563–576.

Westerfield, M. (2000). The Zebrafish Book - A guide for the laboratory use of zebrafish (Danio rerio): University of Oregon Press.

Yablonka-Reuveni, Z., Rudnicki, M. A., Rivera, A. J., Primig, M., Anderson, J. E. and Natanson, P. (1999). The transition from proliferation to differentiation is delayed in satellite cells from mice lacking MyoD. Dev Biol 210, 440–455.

